# Visualizing Liquid Distribution Across Hyphal Networks with Cellular Resolution

**DOI:** 10.1101/2024.08.01.606151

**Authors:** Amelia J. Clark, Emily Masters-Clark, Eleonora Moratto, Pilar Junier, Claire E. Stanley

## Abstract

Filamentous fungi and fungal-like organisms contribute to a wide range of important ecosystem functions. Evidence has shown the movement of liquid across mycelial networks in unsaturated environments, such as soil. However, tools to investigate liquid movement along hyphae at the level of the single cell are still lacking. Microfluidic devices permit the study of fungal and fungal-like organisms with cellular resolution as they can confine hyphae to a single optical plane, which is compatible with microscopy imaging over longer timescales and allows for precise control of the microchannel environment. The aim of this study was to develop a method that enables the visualization and quantification of liquid movement on hyphae of fungal and fungal-like microorganisms. For this, the Fungal-Fungal Interaction (FFI) microfluidic device was modified to allow for the maintenance of unsaturated microchannel conditions. Fluorescein-containing growth medium solidified with agar was used to track liquid transported by hyphae via fluorescence microscopy. Our key findings highlight the suitability of this novel methodology for the visualization of liquid movement by hyphae over varying time scales and the ability to quantify the movement of liquid along hyphae. Furthermore, we showed that at the cellular level, extracellular hyphal liquid movement can be bidirectional and highly dynamic, uncovering a possible link between liquid movement and hyphal growth characteristics. We envisage that this method can be applied to facilitate future research probing the parameters contributing to hyphal liquid movement and is an essential step for studying the phenomenon of fungal highways.

## I. INTRODUCTION

Filamentous fungi and the fungal-like oomycetes are present in a wide range of ecosystems where they perform essential functions^1, 2^. Within soil, filamentous fungi can mediate the cycling of nutrients and are responsible for the decomposition of organic matter^3^. Oomycetes have also been found to contribute to organic matter decomposition in soils^4^. However, the impact of fungi and fungal-like organisms on other important abiotic soil processes, particularly the movement of liquids like water, is not well understood^5^.

The physical structure of soil is composed of a series of solid aggregates surrounded by air pores interconnected by liquid-filled areas, generating irregular regions of liquid saturation and variable nutrient availability^6, 7^. As a result, microorganisms that require a continuous aqueous film for motility, such as bacteria, are unable to mobilize across the air pores in soil^8, 9^. Conversely, the hyphal growth form of filamentous fungi and fungal-like organisms enables them to breach air-liquid boundaries and bridge unsaturated regions into greater volumes of soil and access additional sources of nutrients^10^. Since liquid availability within soil is sporadic, it is thought that hyphae could redistribute water in soil throughout their mycelial network to regions where there is lower water content^5^. This is predominantly believed to be a passive mechanism as hyphae act as hydraulic conductors within soil that provide a path through which liquid can flow along via a concentration gradient^11^. Yet, given that hyphae can modify their surface properties through the production of hydrophobins and other biosurfactants, an ‘active’ component of liquid transport regulated by hyphae is also likely to exist^5, 12, 13^. Understanding the mechanisms of liquid (including water and solubilized organic and inorganic matter) movement by hyphae is important as it may have significant implications for the management of drought^14^. In addition, the ability to transport liquid externally on the hyphal surface could be a key determinant regarding whether fungal and fungal-like species can be used as a “fungal highway” to disperse motile bacteria in soils^15^. Lastly, hyphal-mediated liquid movement might be an essential factor for the successful application of fungal and fungal-like species for bioremediation or enhancing phytoremediation by plants in contaminated soil environments^16, 17^.

An increasing number of studies have demonstrated hyphal-mediated liquid movement experimentally using different fungal and fungal-like species^11, 14, 18-25^. For example, Guhr *et al*. investigated hydraulic redistribution of water by *Agaricus bisporus* using two-chamber mesocosms connected by bridges of hyphae^21^. A three-fold increase in the amount of water redistributed was observed compared to the controls where hyphal bridges between compartments had been cut, in agreement with an identical experiment using *Schizophyllum commune*^14, 21^. Likewise, Worrich *et al*. showed that fungal species *Fusarium oxysporum* and *Lyophyllum* sp. Karsten as well as a fungal-like Oomycete, *Pythium ultimum*, could transport water, carbon, and nitrogen to an environment devoid of these nutrients, which subsequently promoted the germination of bacterial spores^22^. Nevertheless, there are a lack of studies which have visualized and quantified hyphal liquid movement at the cellular level.

Single cell methodologies are fundamental to uncover how microorganisms interact with one another as well as how they interact with their environment at the relevant scale at which these interactions occur^26^. Microfluidic devices are becoming increasingly utilized for studying microorganisms, including fungi and fungal-like organisms, at cellular resolution as they permit precise control of the microchannel environment and can confine microbes to a single optical plane^27^. Typically, microfluidic devices require the microchannels to be filled with liquid medium for inoculation or to maintain the growth of hyphae. However, several studies have begun to develop suitable microfluidic devices for examining the growth of hyphae in partially liquid-filled or air-filled microchannels that more closely simulate the heterogenous liquid distribution found in soil pores^28-31^.

The aim of this study was to develop a method that enables the visualization and quantification of liquid movement by fungal and fungal-like hyphae at the level of the single cell within a microfluidic device. To achieve this, we used a fluorescein-containing growth medium solidified with agar, which could support the growth of fungal-like hyphae and be used to track liquid transported by hyphae via fluorescence microscopy. In this study, we make use of the microfluidic Fungal-Fungal Interaction (FFI) device developed by Gimeno, Stanley *et al*., which was previously established to be a suitable platform for high-resolution time-lapse imaging of hyphae in both liquid-filled and unsaturated (i.e., non-liquid-filled) microchannel conditions^31^. Our key findings highlight the suitability of this novel methodology for enhancing the visualization of liquid films surrounding hyphae via fluorescence microscopy. Microscopy data was quantifiable and demonstrated that hyphal liquid movement is bidirectional and highly dynamic, perhaps closely influenced by hyphal growth and mycelial network structure. This method can be applied to facilitate future research probing the parameters contributing to hyphal liquid movement at the cellular level.

## II. MATERIALS AND METHODS

### A. Strain and Culture Conditions

The fungal-like Oomycete *P. ultimum* M194 was provided by the Laboratory of Microbiology (University of Neuchâtel, Switzerland). Active cultures of *P. ultimum* were maintained on potato dextrose agar (PDA) (Thermo Fisher Scientific) at 26 °C throughout this study. The fluorescein-containing medium used to detect liquid movement across the mycelium was prepared by adding a 66 µM solution of filter-sterilized fluorescein sodium (Sigma-Aldrich) dissolved in Milli-Q water to autoclaved molten PDA in a 10:1 ratio, to give a final concentration of 6.6 µM of fluorescein sodium in the medium. Following this, the fluorescein-containing PDA was poured into sterile Petri dishes (diameter = 90 mm) and allowed to set to form a solid medium. Prior to microfluidic device inoculation, plugs of *P. ultimum* mycelia cultured on PDA were cut using a cork borer (diameter = 4 mm) from the growing edge and placed in the center of either a Petri dish containing PDA or PDA containing fluorescein and grown at 26 °C for 2 to 3 days.

### B. Growth on Media Containing Fluorescein

Plugs containing mycelia were cut from the growing edge of an active culture of *P. ultimum* using a cork borer (diameter = 4 mm) and placed in the center of a Petri dish (diameter = 90 mm) containing either PDA or PDA with fluorescein. Growth was assessed by measuring the radius from the edge of the inoculated agar plug to the edge of the mycelial colony using electronic calipers in two directions to calculate the mean radial growth. Plates were stored at 26 °C and the radial growth of the mycelia was measured for up to 7 days (**Supplementary Material Figure 1**).

### C. Microfluidic Device Fabrication

A detailed protocol for fabricating the FFI microfluidic device has been described previously^31, 32^. In brief, a photolithography mask was used to guide the formation of the device microstructures upon exposing a silicon wafer spin-coated with SU-8 photoresist (MicroChem; final height = 10 µm) to UV light. Afterwards, the photoresist was developed to produce a silicon wafer master mold, and then silanized using chlorotrimethysilane (Merck Life Sciences). Poly(dimethylsiloxane) (PDMS) was prepared by mixing the base with the curing agent in a 10:1 ratio (Sylgard 184 elastomer kit, VWR), which was then degassed under vacuum to remove residual air bubbles. The PDMS was poured onto the master mold and cured at 70 °C overnight. Next, the cured PDMS was removed from the mold, cut into individual slabs and the two inlets were punched using a cutter (SLS; diameter = 4.75 mm). The cut PDMS slabs were washed and dried according to the washing protocol described by Masters-Clark *et al*. prior to bonding onto glass-bottomed Petri dishes (World Precision Instruments; diameter = 50 mm) using a plasma cleaner (Zepto, Diener Electronic)^32^. Bonded FFI devices were stored at room temperature for a minimum of 3 days preceding their use in experiments.

### D. Inoculation of FFI Devices

To maintain humidity, a 100 µL droplet of Milli-Q water was pipetted onto the outer edge of the Petri dish base of the FFI devices. In all experiments, a plug of PDA (4 mm in diameter) was added to the target media inlet of the FFI device. Depending on the nature of the experiment, a plug of media (diameter = 4 mm) with or without mycelia was added to the inoculant inlet. All devices were sealed with parafilm and stored at 26 °C throughout the imaging period for up to 7 days post-inoculation.

### E. Image Analysis and Data Quantification

Phase contrast (PC) and fluorescence microscopy images were obtained using inverted microscopes (Eclipse Ti-U and Eclipse Ti-2, Nikon, **Supplementary Material Table 1**). Image analysis was performed using ImageJ^33^. The ‘rectangle’ tool in ImageJ was used to create a region of interest (ROI) to measure the mean grey value of each diamond within the FFI device in the fluorescence images. To calculate the relative fluorescence intensity of the images obtained daily, the mean grey value of the 1^st^ diamond in the four channels connected to the inoculant inlet was divided by the average mean grey value of the 18^th^ diamond in all four channels connected to the target media inlet combined at each timepoint. To calculate the relative fluorescence intensity for the timelapse images, the mean grey value of diamonds 1 to 6 in the four channels connected to the inoculant inlet were measured then divided by the average mean grey value of the 18^th^ diamond in all four channels connected to the target media inlet combined at each timepoint. To investigate changes in the grey value of hyphae surrounded by liquid films of differing thickness, the ‘line’ tool was used to draw a straight line across hyphae and the pixel intensity across the lines was measured using the ‘plot profile’ analysis tool.

### F. Statistical Analysis

All measured values were inputted into Microsoft Excel (2024), which was used to calculate the mean radial growth of hyphae on media and the relative fluorescence values within the FFI devices for each experiment. In the experiment where microscopy images of the FFI device were taken once per day, data points were removed if there were no hyphae present in the channel or if microchannel quality was insufficient. All subsequent data analysis was performed in RStudio 2023.06.2 with R version 4.2.2, using the tidyverse, dplyr, ggpubr, rstatix, ggsignif, readr and broom packages for statistical analysis and data visualization.

## III. RESULTS

Understanding how fungal and fungal-like hyphae move liquid throughout environments where water and nutrient distribution is heterogenous, such as soil, is of notable importance as it may form a key role within ecosystem functioning^5^. Moreover, little is known about how hyphae influence liquid transport at the microscale, which is the pertinent scale at which hyphae interact with their environment, as well as with other organisms. Microfluidic technologies provide an appropriate platform for investigating hyphal liquid transport due to the ability to control microchannel conditions precisely and confine hyphae to a single focal plane for high-resolution single-cell microscopy imaging.

This study utilizes the FFI microfluidic device, previously developed by Gimeno, Stanley *et al*., which consists of a PDMS slab with embossed microchannels bonded to a glass-bottomed Petri dish that can be sealed with parafilm following inoculation (**Figure 1A**)^31^. There are two inlets that each connect to four microchannels; two of these microchannels connect to both inlets and act as ‘interaction’ channels, whilst two of the microchannels have dead ends and act as ‘control’ channels (**Figure 1B**). Hyphae grow in a single focal plane within the microchannels which are 10 µm in height and the diamond shapes restrict the number of hyphae that can grow into the adjacent diamond at a time. To permit the study of hyphal liquid movement at the level of single cells, it was essential that microchannel conditions could be maintained in an unsaturated state, whilst still sustaining hyphal growth. The FFI device has been established to support the growth of fungal hyphae in microchannels filled with no liquid medium^31^. Further alterations to the device operation were made in this study to ensure microchannel environments were unsaturated by increasing the time following device bonding (i.e., before inoculation) to a minimum of 3 days. This was performed to ensure the plasma-exposed PDMS surfaces had reverted back to a hydrophobic state and that liquid movement into the microchannels was not caused by filling via capillary action^34, 35^.

**Figure 1:**
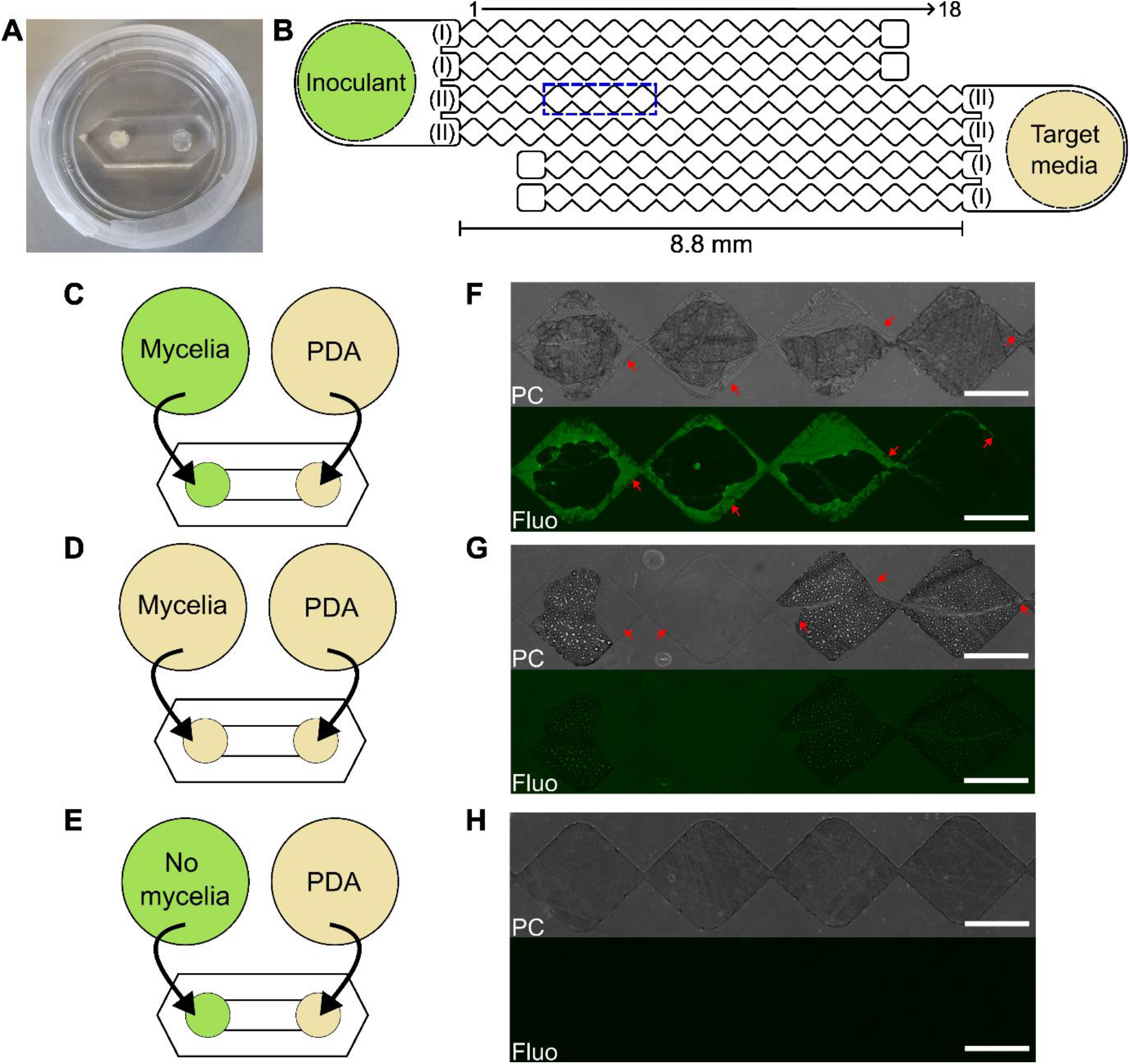
**A**) Image of an inoculated Fungal-Fungal Interaction (FFI) device showing a poly(dimethylsiloxane) (PDMS) slab with embossed microchannels bonded to a 50-mm-diameter glass-bottomed Petri dish. **B**) Schematic of FFI device used in this study and developed by Gimeno, Stanley *et al*.^31^. There are 4 microchannels connected to each inlet, 2 control channels with dead-ends (I) and 2 interaction channels that connect to both inlets (II). Microchannels are formed of 15 to 18 diamond shapes, numbered 1 to 18 from left to right, with each microchannel being 10 µm in height and up to 8.8 mm in length. The dashed blue box corresponds to where the microscopy images in **F-H** were taken on the device. **C-E**) Diagrams illustrating the different conditions tested within the FFI device. In all conditions, a 4-mm-diameter plug of PDA is added to the target media inlet of the FFI device. The inoculant inlet contains a 4-mm-diameter plug of either: *P. ultimum* mycelia grown on PDA containing 6.6 µM of fluorescein (**C**); *P. ultimum* mycelia grown on PDA (**D**); or 6.6 µM of fluorescein sodium with no mycelia (**E**). **F-H**) Representative images of phase contrast (PC) and fluorescence (Fluo) microscopy images of FFI devices at 4 days post-inoculation with *P. ultimum* + fluorescein (**F**), *P. ultimum* only (**G**) and fluorescein only (**H**). Red arrows highlight where liquid is present and scale bars = 250 µm.

### A. Visualization of Hyphal Liquid Transport

Although liquid films surrounding hyphae within unsaturated microchannels can be observed using bright field (BF) and phase contrast (PC) microscopy, it is not always possible to identify the presence of these films. Furthermore, it is often hard to differentiate liquid films moved by hyphae from condensation on microchannel surfaces. The movement of liquid by hyphae is also difficult to quantify as the contrast to the background of the microchannels is poor. To address these issues, we improved visualization of liquid movement by hyphae by incorporating fluorescein into the solidified agar-based culture medium used to grow *P. ultimum* and imaged this using fluorescence microscopy.

Three experimental conditions were set up in this study (*P. ultimum* and fluorescein; *P. ultimum* only; and fluorescein only) (**Figure 1C-E**). In all experimental conditions, a plug of PDA was added to the target media inlet to encourage directional growth of hyphae (**Figure 1C-E**). Whilst the FFI device permits dual inoculation of fungal and fungal-like mycelia, only the left inlet was used as an inoculant inlet in all experimental conditions detailed in this study to reduce the number of variables influencing hyphal liquid transport. The combination of *P. ultimum* and fluorescein was used to test the effectiveness of the method for visualizing hyphal liquid transport and offer a means to quantify it. Since the impact of adding fluorescein sodium to media on mycelial growth is not known, the growth of *P. ultimum* on PDA and the fluorescein-containing media was compared. Statistical analysis revealed that there was no significant difference between the mean radii of the mycelial colonies grown on PDA or PDA containing fluorescein (**Supplementary Material Figure 1**). Hence, the addition of fluorescein to the solid media does not affect hyphal growth. In addition, the *P. ultimum* only condition, in which mycelia were grown on PDA without any fluorescein, served as a control to validate that visualization of hyphal liquid movement is enhanced by the addition of fluorescein (**Figure 1D**). Finally, the fluorescein only condition, whereby a plug of fluorescein-containing media (containing no mycelium) was inoculated into the inlet of the FFI device, was employed as a control to assess whether the fluorescein-containing media could enter the microchannels via capillary action without the presence of hyphae and to identify any potential optical effects from the fluorescent plug or empty microchannels (**Figure 1E**).

Microscopy images of FFI devices were taken once per day at 1, 2, 3, 4 and 7 day(s) post-inoculation for the three experimental conditions indicated above. Examples of phase contrast and fluorescence images of each experimental condition from 4 days post-inoculation are shown in **Figure 1F-H**. Liquid films of varying thickness surrounding or preceding hyphae were visible in phase contrast images of both the *P. ultimum* and fluorescein and the *P. ultimum* only conditions (**Figure 1F-G**). However, the addition of fluorescein to the growth medium in the *P. ultimum* and fluorescein condition unequivocally improves the visualization of these liquid films and is clearly distinguishable from condensation on the microchannel surface (**Figure 1F-G**). No medium is observable within the microchannels in the fluorescein only condition, thus any movement of fluorescein-containing medium into the channels is presumed to be hyphal-mediated (**Figure 1H**).

### B. Quantification of Hyphal Liquid Transport

To quantify hyphal-mediated liquid movement, mean grey values from the fluorescence images were measured to calculate relative fluorescence intensities as detailed in Materials and Methods. To begin with, the relative fluorescence intensity was determined using microscopy images taken once per day to see if there were any statistically significant differences between the relative fluorescence intensities of *P. ultimum* and fluorescein, *P. ultimum* only and fluorescein only conditions and uncover any trends over time (**Figure 2**). As the growth distance of hyphae in each channel was variable, initially only the 1^st^ diamond of each microchannel was used to quantify hyphal liquid movement because this diamond nearly always contained hyphae.

**Figure 2:**
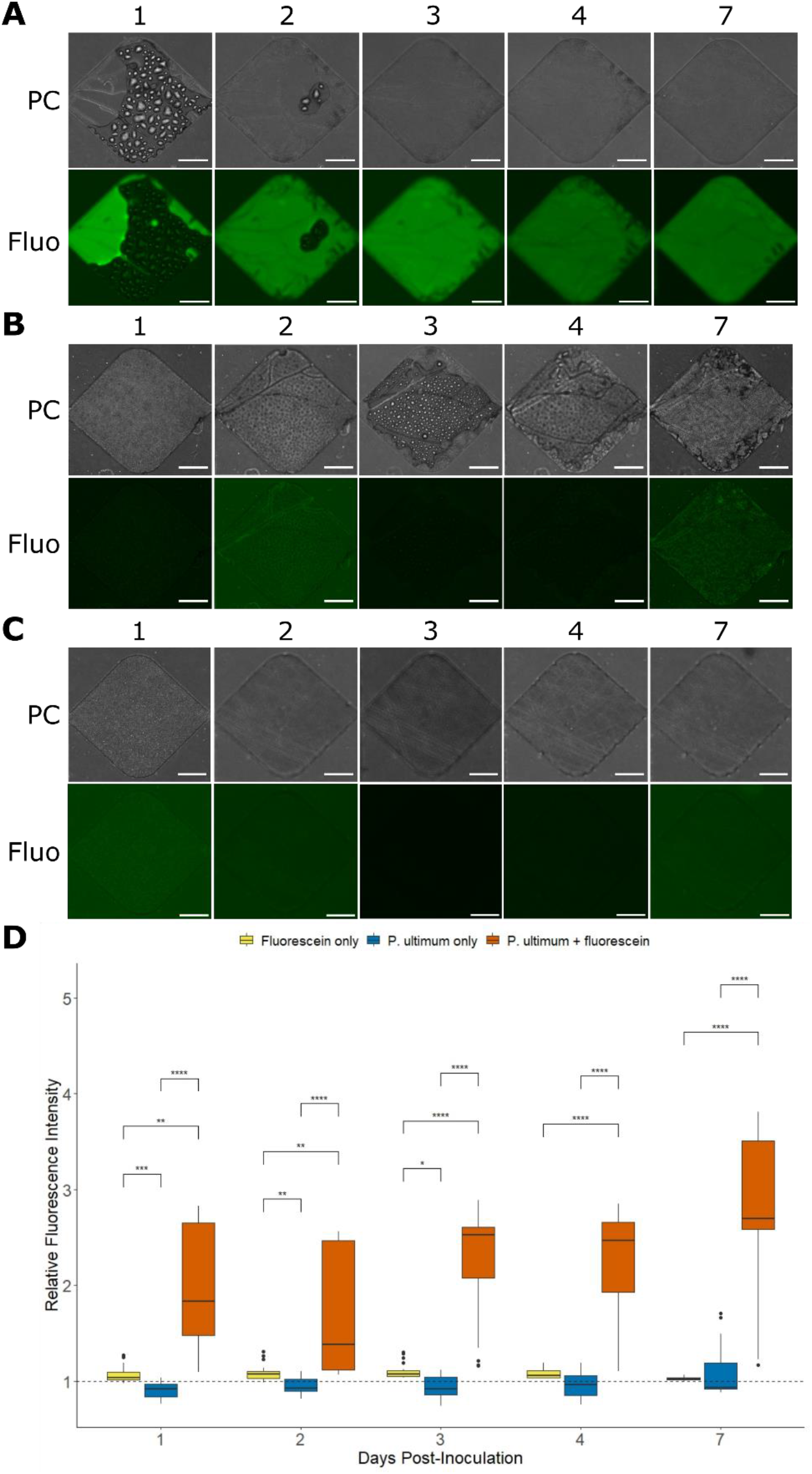
**A-C**) Examples of phase contrast (PC) and fluorescence (Fluo) microscopy images showing the 1^st^ diamond of the Fungal-Fungal Interaction (FFI) device microchannels at 1, 2, 3, 4, and 7 days post-inoculation with *P. ultimum* and fluorescein (**A**), *P. ultimum* only (**B**) and fluorescein only (**C**). Scale bars = 100 µm. **D**) Graph showing the relative fluorescence intensity of the 1^st^ diamond within the microchannels connected to the inoculation inlet, calculated by dividing the mean grey value of the 1^st^ diamond by the average mean grey value of the 18^th^ diamonds measured in the fluorescence images. Dashed line across the plot shows the value at which the relative fluorescence intensity is 1. The box of the plot contains the interquartile range, and the central black line denotes the median, the upper and lower whiskers show the minimum and maximum values and black dots represent outliers. The experiment was repeated 3 times with 2 devices as technical replicates per biological replicate (n ≥ 11). The relative fluorescence intensity of each microchannel was measured and any microchannels in the *P. ultimum* and fluorescein or *P. ultimum* only condition where hyphae did not grow, or any microchannels in all conditions with fabrication issues were removed. A Kruskal-Wallis’s test was performed and found a significant difference (*p* = > 0.05) between the relative fluorescence intensity at each day post-inoculation. This was followed up by a Dunn’s post-hoc test to identify pairwise differences and significance is indicated on the graph by an asterisk (* *p* ≤ 0.05; ** *p* ≤ 0.01; *** *p* ≤ 0.001; **** *p* ≤ 0.0001).

Imaging data from fluorescence microscopy experiments revealed that there was no observable movement of fluorescein into the 1^st^ diamond in the absence of hyphae at any timepoint (**Figure 2C**). Although liquid films surrounding hyphae were evident in phase contrast images of the 1^st^ diamond in both *P. ultimum* and fluorescein and *P. ultimum* only conditions (**Figure 2A-B**), they are only discernible in fluorescence images when both hyphae and fluorescein are present (**Figure 2A and 2D**). In days 1 to 3 post-inoculation, a significant difference was observed between the *P. ultimum* only and fluorescein only controls, which is likely due to condensation in unoccupied microchannel diamonds increasing the background mean grey value, therefore reducing the relative fluorescence intensity in the *P. ultimum* only condition (**Figure 2D**). Importantly, there is a statistically significant difference (*p* ≤ 0.01) between the relative fluorescence intensity of *P. ultimum* and fluorescein and the two control conditions at all timepoints (**Figure 2D**). Thus, the addition of fluorescein to solid agar-based media can help to quantify hyphal-mediated liquid transport within the microchannels of the FFI device as it can be differentiated from condensation present within microchannels in fluorescence microscopy images. Notably, there is no passive filling of microchannels with fluorescein via capillary action in the absence of hyphae.

It was anticipated that the relative fluorescence intensity of the *P. ultimum* and fluorescein condition would steadily increase over time as more hyphae enter the microchannel, transporting more liquid surrounding their surface (**Figure 2A**). Conversely, in the quantified data, the median relative fluorescence intensity decreases at 2 days post-inoculation before increasing again from 3 days post-inoculation (**Figure 2D**). Furthermore, there is large variability in the relative fluorescence intensity of the 1^st^ diamond within the *P. ultimum* and fluorescein condition, which can be assumed to be due to differing amounts of liquid moved by hyphae in each microchannel (**Figure 2D**). Whilst the results from this experiment demonstrate the effectiveness of this method at quantifying hyphal liquid transport, a drawback of taking images once per day is that occurrences at smaller timescales are not observed. In a similar way, because only the relative fluorescence intensity of the 1^st^ diamond was analyzed, if liquid movement is more concentrated at the hyphal front, then this observation may have been missed as hyphae progressed further into the microchannel. Clearly, these results suggest that liquid movement by hyphae may be more dynamic than previously thought and do not simply move along the hyphal surface along a gradient in one direction.

To further demonstrate that the addition of fluorescein to the media can be used to quantify hyphal liquid transport at the level of the single cell, we looked at different hyphae within one region of interest to see if we could exemplify how this method could be applied to identify differences in the thickness of liquid films surrounding hyphae (**Figure 3**). A phase contrast and fluorescence microscopy image from the *P. ultimum* and fluorescein condition at 7 days post-inoculation was selected. Measurements of the grey value were taken by drawing profile lines, ranging from 15.6 to 17.6 µm in length, through four hyphae within the same diamond that were representative of the following hyphal types: I) a hypha surrounded by an easily visible, thicker liquid film; II) a hypha not surrounded by a visible liquid film; III) a hypha surrounded by a poorly visible, thinner liquid film; or IV) a hypha saturated in liquid (**Figure 3A and 3C**). Following this, the grey values every ∼0.65 µm across the lines were plotted (**Figure 3B**).

**Figure 3:**
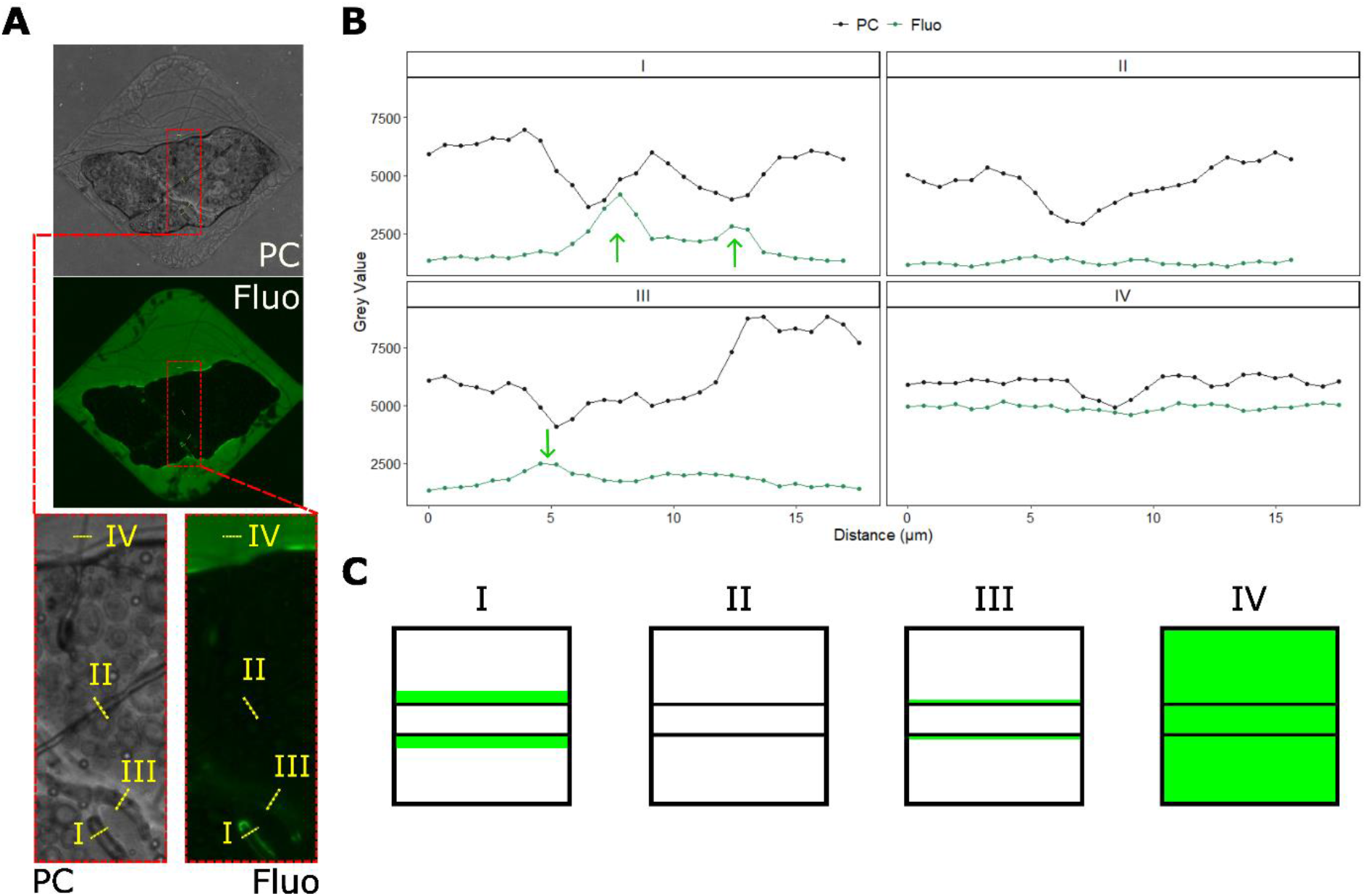
**A**) Phase contrast (PC) and fluorescence (fluo) microscopy image of *P. ultimum* and fluorescein at day 7 post-inoculation. Dashed red boxes and lines indicate close up images of the corresponding larger image. In ImageJ, the ‘line’ tool was used to draw profile lines across hyphae (as indicated by the yellow dashed lines) and subsequently measure the grey value every 0.65 µm. **B**) Graph shows the grey value of each hypha, as labelled in **A** and represented in **C**, over the distance of the measurement line. Green arrows highlight peaks in the grey value of the fluorescence images that correspond to where the liquid film is present. **C**) Schematic illustrating the different hyphae labelled in **A** and measured in **B**. The hyphae were classified as: (I) surrounded by an easily visible, thicker liquid film; (II) not surrounded by a visible liquid film; (III) surrounded by a poorly visible, thinner liquid film; or (IV) saturated in liquid.

In all plots of **Figure 3B**, the grey value of the phase contrast microscopy image drops at the distances surrounding the edges of the hypha. In comparison, there are two clear peaks in the grey value of the fluorescence microscopy image in plot I of the hypha that was surrounded by a thicker liquid film at distances where the liquid film was present around the outside of the hypha. When no visible liquid film surrounding the hypha was visible, there is no peak in the grey value of the fluorescence image in plot II. Likewise, in plot III where a thin liquid film surrounding the hypha was present, there is only one small peak detectable in the grey value of the fluorescence image where the liquid film is marginally thicker on one side of the hypha so can be differentiated from the background. Lastly, in plot IV from the hypha that was saturated in liquid, the grey value over the entire distance of the line in the fluorescence image is higher than the background fluorescence found in the other plots and there are no peaks as the hypha is completely covered by liquid.

### C. Exploring the Dynamics of Hyphal Liquid Transport

To uncover more about the dynamic nature of hyphal liquid transport and further probe the variability of the relative fluorescence intensity of the *P. ultimum* and fluorescein condition, timelapse experiments were conducted to investigate hyphal liquid transport at smaller time increments. Two timelapse experiments using different biological replicates of the *P. ultimum* and fluorescein condition, paired with two timelapse experiments of the fluorescein only control were imaged every 2 hours for 60 hours in total (**Figure 4A, Figure 4B and Supplementary Videos 1-4** (multimedia available online)). Subsequently, the relative fluorescence intensity in the fluorescence images of each of the first 6 diamonds was quantified, as described in the methods, to produce the graphs in **Figure 4C-D**. In both timelapse experiments of the *P. ultimum* and fluorescein condition, examples of bidirectional liquid transport are visible (**Supplementary Videos 1-4, Figure 4A-B**). Visually, differences in the amount and the manner with which liquid was transported by hyphae as well as the way in which the hyphal network developed are apparent between the two timelapse experiments (**Figure 4A-B**). Furthermore, there was a statistically significant difference between the two timelapse experiments of the *P. ultimum* and fluorescein condition in the relative fluorescence intensity of each diamond when all timepoints were combined (**Figure 4C-D**). This finding highlights how the transport of liquid by hyphae is likely linked to hyphal growth and reveals a key application for the presented methodology for exploring how different hyphal properties may influence liquid movement across mycelial networks.

**Figure 4:**
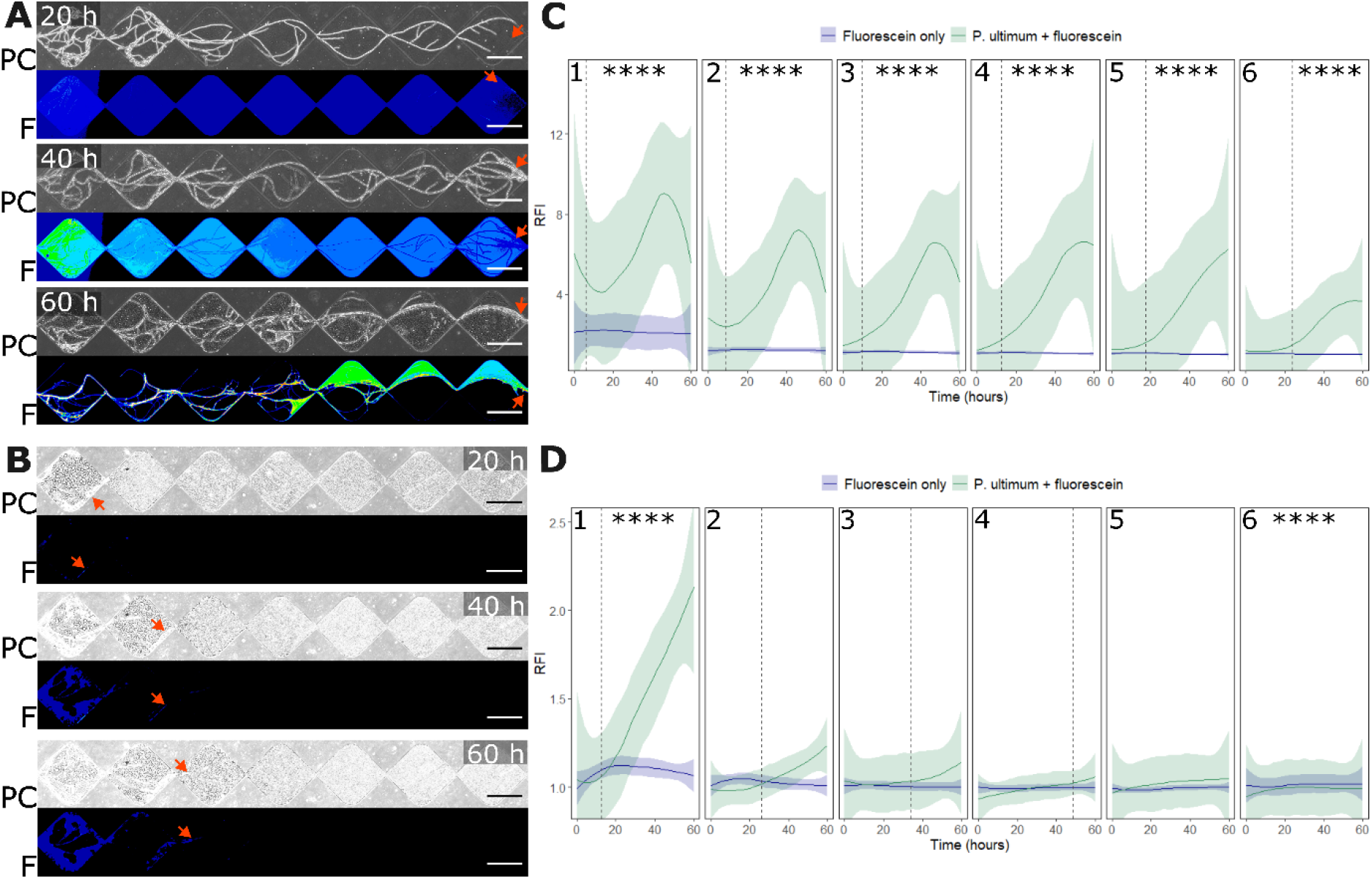
**A-B**) Stills of phase contrast (PC) and fluorescence (F) microscopy images showing the first 6 diamonds of a microchannel of the FFI device at 20, 40 and 60 hours for the *P. ultimum* and fluorescein condition in two different biological replicates. Images were taken every 2 hours for 60 hours and timelapse videos can be found in **Supplementary Videos 1-4** (multimedia available online). Red arrows indicate the front of liquid film and scale bars = 250 µm. **C-D**) Graphs show the relative fluorescence intensity (RFI) of the first 6 diamonds of the *P. ultimum* and fluorescein condition and the fluorescein only condition every 2 hours for a duration of 60 hours. Numbers 1 to 6 at the top of the graphs indicate the diamond number it corresponds to. The RFI was calculated by measuring the mean grey values of diamonds 1 to 6 at each time point in the fluorescence image, then dividing each value by the background mean grey value as described in the methods. The vertical dashed line on the plots shows the mean timepoint when hyphae entered the diamond and lines were only added if hyphae entered the diamond in all four microchannels. Solid blue and green lines represent the mean relative fluorescence intensity and shaded color shows the standard deviation (n = 4). A Wilcoxon signed-rank test was performed and found a significant difference (*p* ≤ 0.05) between the relative fluorescence intensity between the replicates, and between the *P. ultimum* and fluorescein and fluorescein only condition in all diamonds in **C**, and diamonds 1 and 6 in **D**. This was followed up by a Dunn’s post-hoc test to identify pairwise differences and significance is indicated by an asterisk (* *p* ≤ 0.05; ** *p* ≤ 0.01; *** *p* ≤ 0.001; **** *p* ≤ 0.0001).

Certainly, a clear example of the dynamic nature of hyphal liquid transport is exhibited in **Figure 4A**. Microscopy images of the *P. ultimum* and fluorescein condition show the first 6 diamonds of all microchannels are saturated with liquid by 20 hours (**Figure 4A**). Hyphae grew very quickly and entered the 1^st^ diamond in all microchannels by 20 hours (**Figure 4A**). In the *P. ultimum* and fluorescein condition, the relative fluorescence intensity increases substantially in diamonds 1 to 4 until roughly 46 hours where it reaches its peak (**Figure 4C**). Liquid can be observed moving into the microchannels at 20 and 40 hours, mostly in one direction from diamond 1 to the adjoining diamonds further into the microchannel (**Figure 4A**). At 60 hours in the phase contrast microscopy images, hyphae can be seen retracting their cytoplasm and the number of hyphae within the microchannels appears to reduce (**Figure 4A**). Meanwhile, in the fluorescence microscopy images, much of the liquid present in the microchannels at 40 hours disappears by 60 hours, which corresponds to the sudden drop in the relative fluorescence intensity shown in the graph (**Figure 4A and 4C**). At the end of the timelapse at 60 hours, a few films of liquid are still visible in the phase contrast images (**Figure 4A**). Concurrently, in the fluorescence image, fluorescence can be seen surrounding hyphae, with some liquid films present in a few of the diamonds (**Figure 4A**). Statistical analysis of the quantified data for the first replicate confirmed when all timepoints were combined per each diamond that there is a statistically significant difference between the relative fluorescence intensity of the *P. ultimum* and fluorescein and the fluorescein only conditions in all diamonds (**Figure 4C**).

On the other hand, the growth of hyphae in **Figure 4B** was far slower and few instances of hyphal retraction were seen. In the *P. ultimum* and fluorescein condition, the relative fluorescence intensity of the 1^st^ diamond increases from 12 hours, which is also when hyphae begin to grow into the microchannels and enter the 1^st^ diamond at an average of 13 hours (**Figure 4B and 4D**). The volume of liquid in the 1^st^ diamond gradually increases throughout the remainder of the timelapse, entering the 2^nd^ and 3^rd^ diamonds by 40 hours, and hyphae grew into diamond 2 on average at 26 hours and diamond 3 at 34 hours (**Figure 4B and Figure 4D**). Again, the rising volume of liquid entering diamonds 1-3 corresponds to the increasing relative fluorescence intensity shown in the graph (**Figure 4D**). Whilst a few hyphae did reach diamond 4 in all microchannels and diamond 5 in two of the microchannels, only a small increase in the relative fluorescence intensity of the *P. ultimum* and fluorescein condition was detected when compared to the fluorescein only control (**Figure 4B and 4D**). However, when the data for all timepoints is combined, there is a statistically significant difference between the *P. ultimum* and fluorescein condition and the fluorescein only control in the 1^st^ diamond and the 6^th^ diamond (where, in the latter case, the relative fluorescence intensity of the fluorescein only condition is higher than the *P. ultimum* and fluorescein condition) (**Figure 4D**). As the leading hypha did not reach the 6^th^ diamond in all but one microchannel in the *P. ultimum* and fluorescein condition, we postulate that the significance observed here may be due to condensation present in the unoccupied 6^th^ diamond reducing the mean grey value of diamond 6, thus resulting in the background mean grey value used to calculate the relative fluorescence intensity being higher than the mean grey value of diamond 6.

## IV. DISCUSSION

To permit investigations of hyphal liquid movement at the microscale, the microfluidic FFI device was elected as an appropriate platform for this study as it was already confirmed to support the growth of fungal hyphae in unsaturated microchannels. Furthermore, although hyphal-mediated water movement was not quantified, Gimeno, Stanley *et al*. were able to detect the presence of liquid films surrounding the hyphae of fungal species *Trichoderma rossicum* and *Fusarium graminearum* using this device^31^. The fungal-like Oomycete, *P. ultimum*, was selected as the test organism for this study since previous studies observed external liquid films surrounding the hyphae of *P. ultimum* when grown in liquid unsaturated conditions and it has preexisting use in research examining fungal highways^22, 25, 36^. Indeed, when fluorescein was absent from the growth media, liquid films surrounding the hyphae of *P. ultimum* were visible in phase contrast microscopy images; however, the addition of fluorescein enabled liquid moved by hyphae to be easily recognizable and discernible from condensation present on the microchannel surface. In addition, it was shown that by measuring the grey value of a line across a hypha, the thickness - or absence - of the liquid film was distinguishable. Though *P. ultimum* is classified as an Oomycete, it can be assumed that this method would be applicable to exploring hyphal liquid transport in fungal species as *Pythium* species are known to obtain nutrients from their environment in a comparable way to true fungi^37^. Most previous work examining hyphal-mediated liquid transport has been performed using arbuscular mycorrhizal fungi (AMF), but several studies have confirmed hyphal liquid movement using filamentous Basidiomycota and Ascomycota^14,21,22,25,30^. Nonetheless, it remains unclear whether all fungal and fungal-like species are capable of transporting liquid along hyphae.

The controls used in this study verified fluorescein did not enter the microchannels via capillary action when no hyphae were present. Occasional optical effects from condensation within unoccupied microchannels contributed to outliers in the control conditions. In future studies, altering the microchannel surfaces to PDMS bonded to PDMS, as opposed to PDMS bonded to glass, may help to reduce condensation within the microchannels. We chose to use fluorescein in this study as it is a well-established fluorescent tracer, is highly soluble in water and has low toxicity^38, 39^. Our results found that the addition of fluorescein to PDA has no impact on the growth of mycelia. Liquid transported by hyphae is thought to either travel externally along the cell wall or internally inside the cell membrane^5^. Throughout this study, liquid films along the external surface of hyphae were evident and it was assumed that fluorescence from hyphae came mostly from liquid surrounding the hyphae externally. This is in agreement with another study that showed water was transported externally by AMF hyphae using a fluorescent tracer and heavy water^24^. Nevertheless, the possibility that some instances of internal liquid transport may have been detected during this study cannot be ruled out.

High variability in hyphal liquid transport between biological replicates was found in this study both visually, in the amount of liquid visible in microscopy images, and numerically, in differing relative fluorescence intensities. Whilst probing the factors that influence hyphal liquid transport was beyond the scope of this research, novel insights into how hyphal characteristics may contribute to this were apparent in our findings. Differences in the number of hyphae within microchannels, the speed at which hyphae grew and the thickness and branching behavior of hyphae were all observable between replicates, potentially playing a role in the variable liquid movement. Future research would benefit from examining hyphal liquid movement using a fluorescently tagged fungal or fungal-like species as this would permit the correlation of hyphal growth with liquid movement via fluorescence microscopy. One of the most important findings of this study concerns the fact that hyphal liquid movement is far more dynamic at the cellular level than previously thought, which studies conducted at colony level are not able to detect.

Climate change is estimated to trigger a surge in the frequency and severity of drought worldwide^40^. In response to this, approaches applying inoculants of fungi have been suggested to maintain and enhance soil structure undergoing drought conditions to alleviate the impact of climate change^41^. Hence, single cell investigations of hyphal-mediated liquid transport are vital for improving our understanding of the mechanisms behind liquid movement throughout soil environments to ensure future climate change strategies are ecologically informed. Exploring the ability of hyphae to move liquids across their network could assess the suitability of fungi and fungal-like organisms in bioremediation to remove solubilized contaminants from soils. Baranger *et al*. utilized a microfluidic device to show that the fungus *Talaromyces helicus* could uptake the contaminant benzo(α)pyrene internally via direct contact, and possibly extracellularly along the hyphal surface^42^. Likewise, the liquid films surrounding hyphae can serve as a fungal highway in which motile bacterial species can disperse throughout unsaturated regions of soil, thus enhancing the reach of pollutant-degrading bacterial species for bioremediation^15, 43^. A key determinant of whether hyphae can be used as a fungal highway by bacteria is the thickness of the liquid film surrounding hyphae, which needs to be at least 2 to 10 µm thick^44^. As shown by the findings of this study, the presented methodology can differentiate between the thickness or absence of liquid films surrounding hyphae and highlights a prospective application for screening fungal and fungal-like species for their ability to act as a fungal highway.

## V. CONCLUSIONS

This study aimed to develop a method which could visualize liquid movement by fungal and fungal-like hyphae at cellular level in a way that was quantifiable. By combining the use of the microfluidic FFI device with a fluorescein-containing growth medium, liquid films transported by hyphae were clearly detectable via fluorescence microscopy. Moreover, the microscopy data could be quantified to observe trends in liquid movement over time and differentiate between hyphae surrounded by liquid films of various thickness. Statistically significant differences were found between the relative fluorescence intensity of microchannels from devices inoculated with *P. ultimum* grown on fluorescein-containing media, compared to the controls where *P. ultimum* was grown without fluorescein, or where fluorescein-containing media was present but without mycelia. Notably, the dynamic nature of liquid movement by hyphae was demonstrated and shown to be highly variable between biological replicates, thus liquid transport by hyphae is likely correlated to hyphal growth and network development. In summary, the methodology presented in this study offers many research opportunities for expanding our knowledge of what biotic and abiotic parameters influence hyphal liquid transport in unsaturated environments. In the future, we envision that this methodology could be applied to probe the parameters of hyphal-mediated water transport, such as whether it is a species-specific trait; the extent to which physiochemical surface properties impact hyphal water movement; and whether this phenomenon is due to mass flow or a controllable bidirectional ability.

## SUPPLEMENTARY MATERIAL

Supplementary materials provided include the mean radial growth of *Pythium ultimum* on potato dextrose agar (PDA) and PDA containing fluorescein; a table highlighting the equipment and setting used to obtain microscopy images; and descriptions of each supplementary video. Supplementary videos are available online.

## ACKNOWLEDGMENTS

We acknowledge financial support from the Department of Bioengineering at Imperial College London and The Leverhulme Trust (Research Grant Reference: RPG-2020-352).

## Author Contributions

**Amelia J. Clark:** Conceptualization (supporting), Data curation (lead), Formal analysis (lead), Investigation (lead), Methodology (lead), Validation (lead), Visualization (equal), Writing - original draft (lead). **Emily Masters-Clark**: Conceptualization (supporting), Formal analysis (supporting), Investigation (supporting), Methodology (supporting), Visualization (equal), Writing - review & editing (supporting). **Eleonora Moratto**: Data curation (supporting), Formal analysis (supporting), Visualization (equal), Writing - review & editing (supporting). **Pilar Junier**: Conceptualization (supporting), Funding acquisition (supporting), Resources (supporting), Supervision (supporting), Writing - review & editing (supporting). **Claire E. Stanley**: Conceptualization (lead), Funding acquisition (lead), Methodology (supporting), Project administration (lead), Resources (lead), Supervision (lead), Visualization (equal), Writing - original draft (supporting), Writing - review & editing (lead).

## DATA AVAILABILITY

The data that support the findings of this study are available from the corresponding author upon reasonable request.

