## Supplementary Information for "Visualizing Liquid Distribution Across Hyphal Networks with Cellular Resolution"

### SUPPLEMENTARY MATERIAL

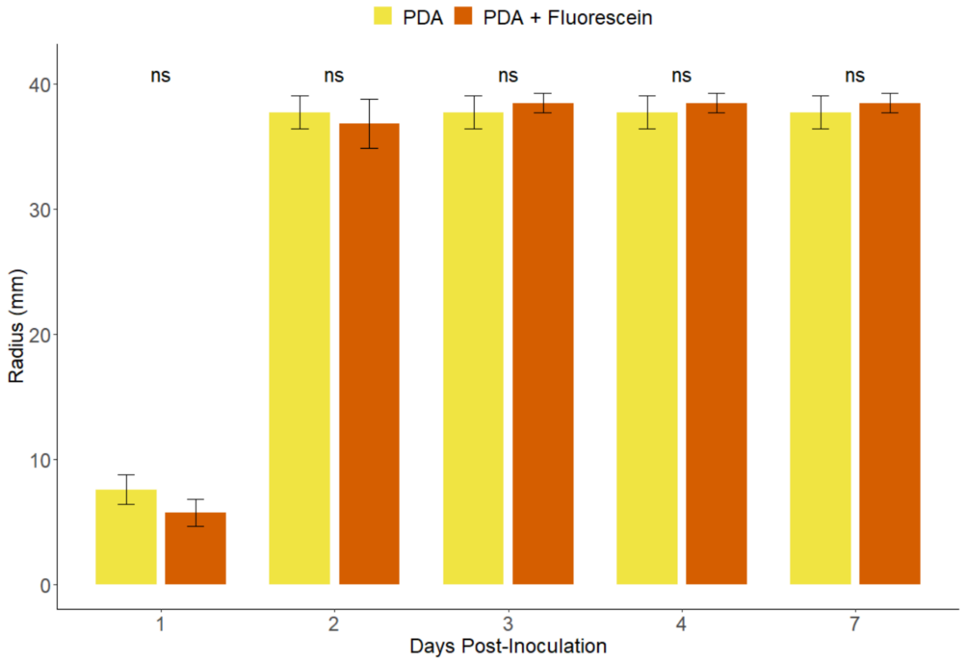


**Supplementary Materials Figure 1**: Graph showing the radial growth of *Pythium ultimum* mycelia on Petri dishes (Ø = 90 mm) containing potato dextrose agar (PDA) or PDA containing fluorescein for up to 7 days post-inoculation. Growth was assessed by measuring the radius from the edge of the inoculated agar plug to the edge of the mycelial colony using electronic calipers in two directions to calculate the mean radial growth (n = 6). To test whether there was a statistically significant difference between the radii of the mycelia grown on PDA alone and PDA with fluorescein, a T-test was performed and found no significant difference, denoted on the graph by ‘ns’. Error bars represent the standard deviation of the mean.

**Supplementary Materials Table 1:** Microscopy Equipment and Settings

| Equipment | Eclipse Ti-U (Nikon) | Eclipse Ti2-E (Nikon) |
|---|---|---|
| Objective | Air immersed x10/0.3 NA Plan Fluor (Nikon) | Air immersed x10/0.3 NA CFI Plan Fluor (Nikon) |
| Camera | Retiga R1 CCD camera (Qimaging) | DS-Qi2 Mono Digital Microscope Camera (Nikon) |
| Stage | Prior Scan III motorized stage | Motorized stage (Nikon) |
| Illumination | High-power light emitting diode (LED) (Omicron-Laserage Laserprodukte GmbH) | High power LED (Nikon) |
| Filter Sets | Band pass filter: 465 nm (Omicron-Laserage Laserprodukte GmbH)<br>Beam splitters: 495 nm (AHF Analysentechnik AG)<br>Emission filters: 525/50 nm (AHF Analysentechnik AG) | GFP-4050B Filter Cube<br>Excitation: EX 466/40<br>Dichroic Mirror DM 495<br>Barrier Filter: BA 525/50 (Nikon) |

Table contains equipment used for obtaining microscopy images. Images obtained once per day (**Figure 1-3**) were taken using the Ti-U microscope and timelapse images (**Figure 4 and Supplementary Videos 1-4**) were taken using the Ti-2 microscope.

**Supplementary Videos 1-4** (multimedia available online)

**Supplementary Video 1:** Timelapse Video of Channel 1 of the FFI Device Inoculated with *Pythium ultimum* in the *P. ultimum* and Fluorescein Condition (Replicate 1).

Time-lapse experiment was recorded over 60 hours with 2 hour intervals between image acquisition using phase contrast (top) and fluorescence (bottom) microscopy. *P. ultimum* can be seen growing across the first 6 diamonds of a channel of the FFI device from the left inoculant inlet. Liquid films are visible surrounding hyphae and are shown to be transported in multiple directions across the mycelial network. 16-color look-up table (LUT) has been applied to the fluorescence timelapse (bottom), scale bars = 250 μm and timestamps are displayed in hours.

**Supplementary Video 2:** Timelapse Video of Channel 4 of the FFI Device Inoculated with *Pythium ultimum* in the *P. ultimum* and Fluorescein Condition (Replicate 1).

Time-lapse experiment was recorded over 60 hours with 2 hour intervals between image acquisition using phase contrast (top) and fluorescence (bottom) microscopy. *P. ultimum* can be seen growing across the first 6 diamonds of a channel of the FFI device from the left inoculant inlet. Liquid films are visible surrounding hyphae and are shown to be transported in multiple directions across the mycelial network. 16-color look-up table (LUT) has been applied to the fluorescence timelapse (bottom), scale bars = 250 μm and timestamps are displayed in hours.

**Supplementary Video 3:** Timelapse Video of Channel 3 of the FFI Device Inoculated with *Pythium ultimum* in the *P. ultimum* and Fluorescein Condition (Replicate 2).

Time-lapse experiment was recorded over 60 hours with 2 hour intervals between image acquisition using phase contrast (top) and fluorescence (bottom) microscopy. *P. ultimum* can be seen growing across the first 6 diamonds of a channel of the FFI device from the left inoculant inlet. Liquid films are visible surrounding hyphae and are shown to be transported in multiple directions across the mycelial network. 16-color look-up table (LUT) has been applied to the fluorescence timelapse (bottom), scale bars = 250 μm and timestamps are displayed in hours.

**Supplementary Video 4:** Timelapse Video of Channel 4 of the FFI Device Inoculated with *Pythium ultimum* in the *P. ultimum* and Fluorescein Condition (Replicate 2).

Time-lapse experiment was recorded over 60 hours with 2 hour intervals between image acquisition using phase contrast (top) and fluorescence (bottom) microscopy. *P. ultimum* can be seen growing across the first 6 diamonds of a channel of the FFI device from the left inoculant inlet. Liquid films are visible surrounding hyphae and are shown to be transported in multiple directions across the mycelial network. 16-color look-up table (LUT) has been applied to the fluorescence timelapse (bottom), scale bars = 250 μm and timestamps are displayed in hours.
